# Validity of Low-Dimensional Representation of Whole-Body Movements

**DOI:** 10.64898/2026.08.31.746368

**Authors:** M. Wakabayashi, A. Katayama, T. Asai

## Abstract

Research on impression evaluation from whole-body movements has traditionally focused on the relationships between emotional perception or aesthetic evaluation and movement patterns characterized by local kinematic features and inter-limb coordination. However, because observers perceive the entire body simultaneously when evaluating emotions or aesthetics in real-world viewing situations, impressions derived from whole-body movements should also be investigated. This requires a low-dimensional representation that summarizes whole-body information in an interpretable manner. In the present study, we evaluated whether the state space generated by multidimensional scaling (MDS) provides a valid representation for summarizing whole-body movement. Specifically, using exercises from *Radio Calisthenics No. 1*, we examined whether distances between points in the MDS state space primarily reflected exercise category rather than individual differences. The results showed that distances in the state space reflected exercise category more strongly than performer identity, suggesting that MDS captures similarities among postures of the same exercise. However, the patterns of distances between conditions varied across exercise categories. These findings suggest that MDS-based state-space representations provide a valid and interpretable approach for summarizing whole-body movements. Nevertheless, further validation using a wider variety of whole-body movements and a more diverse participant population is required.

## 1. Introduction

Humans form subjective impression judgments, such as emotion recognition and aesthetic evaluation, from a wide variety of movements encountered in everyday life. Although emotions are typically conveyed through a combination of verbal, facial, and bodily cues, observers can infer others’ emotional states from movement alone, including gait patterns (Dittrich et al., 1996; Halovic & Kroos, 2018a,b). In the context of aesthetic evaluation, dance has been studied within empirical aesthetics as a dynamic art form, and movement itself has been recognized as one of its fundamental components (Christensen & Calvo-Merino, 2013).

Even under highly restricted conditions in which only body movement is presented through point-light displays representing the major joints of the body, humans can readily perceive biological motion (Johansson, 1973). Furthermore, observers can extract information not only about movement itself but also about the emotions expressed through the movement (Atkinson et al., 2004; Halovic & Kroos, 2018a). In the context of aesthetic evaluation, previous studies have likewise isolated movement information by presenting dance stimuli as stick avatars and demonstrated associations between kinematic parameters and aesthetic evaluations (Torrents et al., 2013). These findings suggest that movement information itself contributes to impression judgments.

A direct relationship between movement kinematics and impression judgments, such as emotion recognition and aesthetic evaluation, has been reported in previous studies (Orlandi et al., 2020; Torrents et al., 2013; Pollick et al., 2001; Sawada et al., 2003). At the same time, kinematic parameters are often complex and highly dependent on the type of movement being performed (Orlandi et al., 2020; Barliya et al., 2013). For example, happy and angry gait patterns are characterized by increased arm movements, longer stride lengths, and faster walking speeds, whereas sad gait patterns tend to involve reduced arm movements, shorter strides, and slower walking speeds (Halovic & Kroos, 2018b). In the context of aesthetic evaluation, studies of leg-raising movements in classical ballet and contemporary dance have shown that aesthetic ratings are associated with variables such as the time required to reach a target posture, leg angle, the perpendicularity of the supporting leg relative to the floor, and lower jerk values (Torrents et al., 2013; Bronner & Shippen, 2015). These findings suggest that both the body regions considered important and the ways in which movement characteristics are quantified vary substantially across studies.

Although quantifying movement characteristics of individual body parts is important for understanding different types of movement, whole-body movements are inherently formed through the coordinated actions of multiple body segments. For example, raising one arm alters the positions of the shoulder and neck, whereas taking a step forward changes the balance of the upper body. Therefore, studies of whole-body movement must take coordination among body parts into account. Otherwise, it becomes difficult to determine whether the characteristics of a specific body part influence impression judgments independently or whether they are associated with impression judgments because they co-vary with other movement features. In studies of gait, coordination among body parts has been shown to influence emotion recognition, including coordinated changes among lower-limb segments and between arm and leg movements (Barliya et al., 2013; Wakabayashi et al., 2026). However, because impression judgments in real-world viewing situations are based on the simultaneous observation of the entire body, it is necessary to investigate the relationship between whole-body posture and impression judgments beyond coordination among specific body parts. Furthermore, considering the whole body simultaneously may allow researchers to examine whether changes in specific body parts or inter-limb coordination influence impression judgments independently of global movement changes or instead arise as a consequence of changes in whole-body movement. Such an approach may deepen our understanding of previously reported relationships between impression judgments and specific body regions or patterns of coordination.

A major challenge in investigating the relationship between whole-body posture and impression judgments without focusing on specific body parts is how to represent complex postural information consisting of multiple kinematic variables in an interpretable form. Because whole-body movement consists of numerous kinematic features that change in a coordinated manner, quantitatively describing its relationship with impression judgments requires a representation of the relationships among postures at individual frames.

However, a variety of dimensionality-reduction approaches have been used to summarize whole-body movement, and no standard method has yet been established. Therefore, the present study aimed to examine an approach for representing whole-body movement in a low-dimensional space in an interpretable manner, with the goal of summarizing complex whole-body movement. Establishing such a representation would provide a foundation for investigating the relationship between whole-body movement and impression judgments.

Beyond serving as a foundation for investigating the relationship between whole-body movement and impression judgments, such a representation may also contribute to a broader understanding of whole-body movement itself. Because whole-body movement is characterized by coordinated interactions among multiple body segments, identifying these coordination patterns requires data-driven exploration based on whole-body features. If the proposed approach proves to be valid, the correspondence between the dimensions of the resulting state space and characteristic postural patterns may reveal which movement patterns constitute independent components of whole-body movement. Furthermore, such a representation may facilitate comparisons across different genres of whole-body movement. Previous studies have typically investigated the kinematic characteristics of movements, such as dance, within individual genres. In contrast, by representing postural configurations in a common low-dimensional space independently of their cultural background or historical development, the proposed approach may enable posture relationships across different movement genres to be described in a directly comparable manner.

Various approaches have been proposed for reducing the dimensionality of movement data. For example, Troje (2002) extracted average postures from gait cycles and reconstructed walking movements using a small number of movement components derived from principal component analysis (PCA). Bekemeier et al. (2019) similarly characterized handover movements in a low-dimensional space using hand kinematics estimated from motion-capture data. Because impression judgments are likely to be influenced by the relative similarities and differences among postures, describing the similarities and differences between postures may provide a useful framework for investigating such judgments. Accordingly, the present study applied MDS. MDS enables multivariate data to be represented in a low-dimensional space on the basis of similarity relationships and has been applied to complex datasets such as whole-brain activity patterns (Asai et al., 2023, 2026).

To evaluate whether whole-body movements could be represented in an interpretable low-dimensional space, we established the following criteria. We conceptualized movement as a sequence of postures and represented each posture at an individual frame as a single point within a state space. Under this framework, a movement can be described as a trajectory of points that changes over time. We then examined whether the resulting representation satisfied two conditions. First, similarities among postures should be reflected in the distances between points in the state space, rather than collapsing into a single cluster or being distributed at uniform intervals. Second, points belonging to the same exercise category should be located closer to one another regardless of the individual performing the exercise or the position and orientation of the body. Because the ultimate goal of this research is to investigate impression judgments, priority was given to distinguishing exercise categories rather than identifying individuals.

A challenge in constructing a state space using MDS lies in the quantity and variety of postures included in the input data. Because MDS represents data according to the relationships among the observations used as input, the resulting state space depends heavily on the range of postures included in the dataset. Therefore, a wide variety of postures was required to construct a meaningful representation of whole-body movement. Traditionally, obtaining accurate movement data has required specialized equipment and experimental environments, often placing a considerable burden on participants (Roggio et al., 2024). In the present study, markerless motion capture was employed to reduce participant burden while enabling repeated short recording sessions across multiple days.

In addition to the recording method, selecting an appropriate movement task was important for efficiently capturing a wide variety of whole-body postures. Because the study required both low participant burden and the performance of the same exercise categories across multiple participants, Radio Calisthenics No. 1 was selected as the experimental task. This exercise is familiar to many Japanese people and is accompanied by standardized demonstration videos. As a result, participants were able to perform the same exercises at similar tempos and timings while engaging the entire body.

## 2. Method

### 2.1 Participants

Five participants (M = 37.4 years, SD = 8.6 years; 3 males and 2 females) took part in the experiment. Each participant completed one recording session per day. Data collection was conducted across one to three days depending on the participant. One participant completed one recording session, three participants completed two recording sessions, and one participant completed three recording sessions.

### 2.2 Procedure

Participants performed Radio Calisthenics No. 1 while watching an instructional video (Radio Calisthenics Channel [Japan Post Insurance Official], 2021). The video was presented on either a laptop computer (MacBook Pro, 13-inch display) or a tablet device (iPad Pro, 11-inch display). The display device was positioned in advance such that the participant’s entire body could be captured by the motion-capture cameras. Participants were instructed to stand in a position from which they could view the instructional video directly and to begin the exercise from that position. After motion-capture recording had started, the instructional video was played, and the experiment began.

Participants’ movements were recorded using seven to eight markerless motion-capture cameras (camera: DSC-RX0 MII, Sony; camera control box: CCB-WD1, Sony). The number of cameras used varied between recording sessions because of differences in camera connectivity. Motion-capture data were recorded at 60 fps. For some recording sessions, an additional video camera was used for synchronization checks; however, these recordings were not included in the analyses.

### 2.3 Data preprocessing

Coordinate estimation was performed after each recording session using Theia3D (Theia Markerless Inc., Kingston, ON, Canada). Because a 20-Hz cutoff was applied during coordinate estimation in Theia3D, no additional smoothing was applied to the exported coordinates. One trial from one participant was excluded from all analyses following visual inspection because camera synchronization and coordinate estimation had clearly failed. In addition, for the participant who completed three recording sessions, one trial showing larger coordinate fluctuations was excluded following visual inspection to reduce imbalance in the number of trials across participants. Consequently, a total of eight trials from five participants were included in the analyses.

Theia3D provided three coordinate points for each of 20 body segments, corresponding to the proximal, center, and distal locations of each segment. Fifteen joint-center points (neck, bilateral shoulders, elbows, wrists, hips, knees, ankles, and feet) were calculated by averaging the proximal and distal coordinates of adjacent body segments. Five terminal points (head, bilateral fingertips, and bilateral toe tips) were represented using the distal coordinates because no adjacent segment was available. For the waist, the segment-center coordinate was used directly. Consequently, a total of 21 body points were used in the subsequent analyses.

Next, preprocessing was conducted to generate the input data for MDS. First, frames containing missing values were removed. Specifically, any frame containing a missing value in at least one of the 21 body points was excluded from further analyses.

Subsequently, the start and end points of each trial were temporally aligned. Following coordinate estimation in Theia3D, the demonstration video visible in the synchronized camera recordings was used as a temporal reference. Specific time points in the demonstration video were defined as the start and end points, and the corresponding frames were identified manually for each trial.

Because markerless motion capture may include estimation errors relative to actual body positions, an additional outlier-removal procedure was applied. For each trial, exclusion was based on the mean displacement across all 21 body points. Consecutive segments containing two or more frames in which the mean inter-frame displacement exceeded the trial mean by more than four standard deviations were removed.

To reduce computational costs during correlation matrix construction and MDS computation, and to improve the interpretability of the resulting state space, temporal downsampling was performed. Recordings were originally captured at 60 fps to ensure visually smooth motion; however, one frame was retained every ten frames, resulting in a sampling rate of approximately 6 fps.

Finally, the coordinate data were transformed into distance vectors referenced to the waist point. Using raw coordinates could cause differences in participants’ standing positions and slight variations in body orientation to influence distances within the MDS state space. Because the purpose of the study was to characterize body posture independently of global position and orientation, Euclidean distances from the waist to each of the remaining 20 body points were calculated for every frame. Each frame was therefore represented as a 20-dimensional vector consisting of these distances. Similarity between frames was quantified using Pearson’s correlation coefficient between the corresponding distance vectors. For MDS, a dissimilarity measure was calculated as 1 - r, where r denotes the correlation coefficient between two postures.

Classical metric MDS was performed using the cmdscale function in R (version 4.5.0; R Foundation for Statistical Computing, Vienna, Austria), and all frames from all trials were embedded into a common three-dimensional state space.

### 2.4 Analysis method

Statistical analyses were conducted using the coordinates obtained from the MDS state space. The primary focus of the present study was not the absolute coordinate locations of points within the state space, but rather the distances between points. To calculate distances within the MDS state space and classify them according to experimental conditions, the Radio Calisthenics sequence was first segmented into individual exercises.

Based on the instructions and transitions between exercises in the demonstration video, the entire sequence was segmented into 13 exercises and subsequently organized into 12 exercise categories by combining exercises of the same type (Figure 1). Next, the exercise transition points in the demonstration video visible in one trial were identified visually, and the onset of each exercise was recorded at the frame level. For the remaining trials, only the start and end points of the entire exercise sequence were identified from the demonstration video. The onset of each exercise was then calculated based on the number of frames for each exercise obtained from the trial in which all exercise onset timings had been recorded. The recordings were segmented such that the duration and timing of each exercise were aligned across all trials.

**Figure 1.**
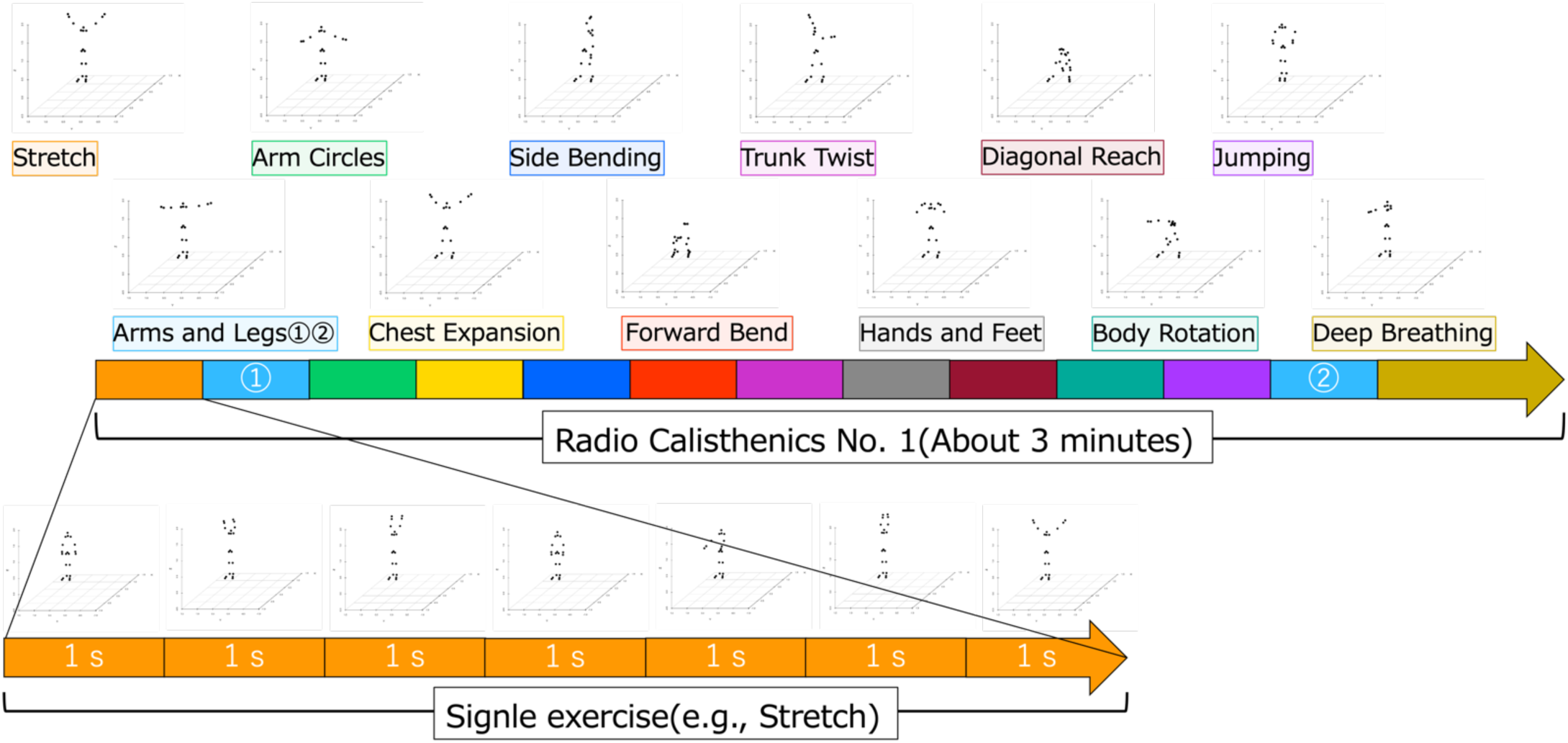
Schematic illustration of the segmentation of Radio Calisthenics. The labels and the colored segments within the arrow are shown in matching colors, indicating the order in which the exercises were presented. The numbered labels indicate the first and second repetitions of the same exercise. The images shown here were created from test recordings collected by the experimenter and were not included in the analysis.

In Radio Calisthenics, a single exercise typically consists of a short action repeated two to eight times. Because each exercise passes through multiple characteristic postures, each exercise was further divided into 1-s bins. For each bin, the centroid of the corresponding coordinates in the MDS state space was calculated and used in subsequent analyses. To control for variability in the number of frames contributing to each bin centroid, bins containing fewer than three frames were excluded from analysis.

Within the MDS state space, all pairwise distances between bin centroids were calculated, with each participant pair treated as a single sample. Distances were classified into four conditions based on the combination of two factors: a person factor indicating whether the exercises were performed by the same or different individuals, and an exercise factor indicating whether the compared exercises belonged to the same or different exercise categories. A mixed-design two-way ANOVA was conducted with the person factor as a between-subject factor and the exercise factor as a within-subject factor. Subsequently, a three-way ANOVA including exercise category as an additional factor was performed. Because distances were aggregated for each exercise category according to whether the compared exercises belonged to the same category as the target exercise, some distance data were included in multiple exercise-category aggregates.

Finally, in a separate three-way ANOVA examining the effects of recording sessions, the exercise-category factor was replaced with a recording-day factor. Because data collection was conducted across four separate recording days, distances were classified according to whether the compared recordings were obtained on the same or different days. As the purpose of this analysis was to examine differences attributable to recording day itself, all available data were included regardless of the number of trials contributed by each participant.

## 3. Results

### 3.1 MDS State Space

First, the correlation matrix used to construct the MDS state space and the resulting frame-wise points within the MDS state space are shown in Figures 2a and 2b, respectively. The correlation matrix obtained by combining all eight trials revealed that correlation coefficients were broadly distributed across both positive and negative values, both within and between trials. The points in the MDS state space were not concentrated around the center but instead formed a roughly spherical distribution. Furthermore, when color-coded by trial, no apparent clustering by participant or trial was observed, and points from different participants and trials were distributed in an overlapping manner.

**Figure 2.**
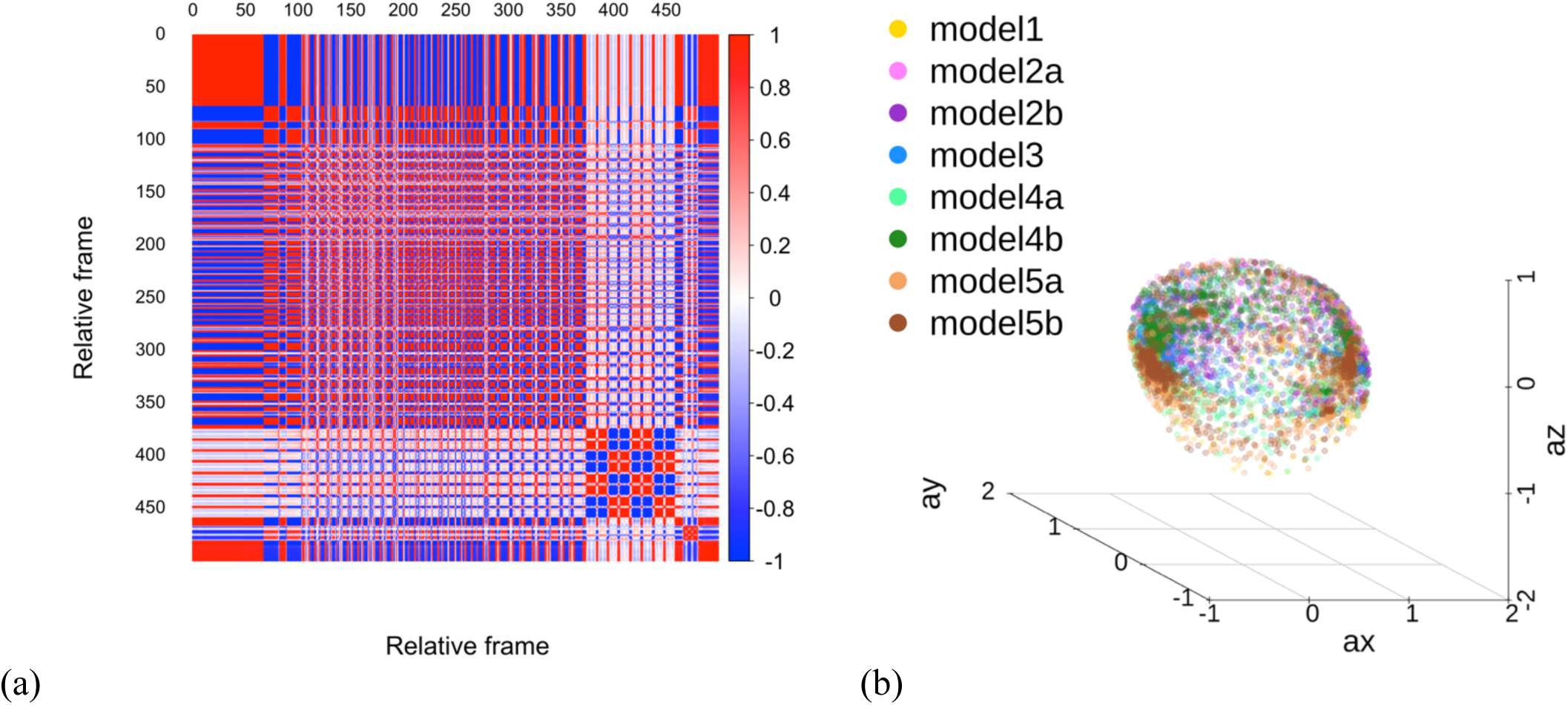
Correlation matrix between postures and plot of all postures in the MDS state space. **(a)** Heatmap of the correlation matrix showing the correlation coefficients between postures, which formed the basis of the distance matrix used as input to MDS. Colors were assigned such that coefficients close to 1 are shown in red, coefficients close to −1 are shown in blue, and coefficients close to 0 are shown in white. The distance matrix used as input to MDS was calculated as 1 − r for each pair of postures. For clarity, only the first 500 frames of the correlation matrix are shown. **(b)** Points representing the postures included in the analysis in the MDS state space. Colors indicate individual trials. Multiple trials from the same participant are represented using similar colors: Participant 1, yellow; Participant 2, light purple and purple; Participant 3, blue; Participant 4, light green and green; and Participant 5, orange and dark brown.

To examine whether different exercise categories showed characteristic distributions within the MDS state space, Figure 3 was created based on the points shown in Figure 2b. Bin-averaged coordinates were superimposed on the frame-wise points and color-coded according to exercise category. Averaging coordinates within each bin resulted in the distribution of bin centroids within the interior of the spherical structure. Nevertheless, some exercises exhibited broader distributions along specific dimensions of the state space. Visually, the exercises Arms and Legs, Chest Expansion, and Deep Breathing were broadly distributed along the first dimension, whereas Side Bending showed a broader distribution along the second dimension. No clear exercise-specific pattern was observed along the third dimension.

**Figure 3.**
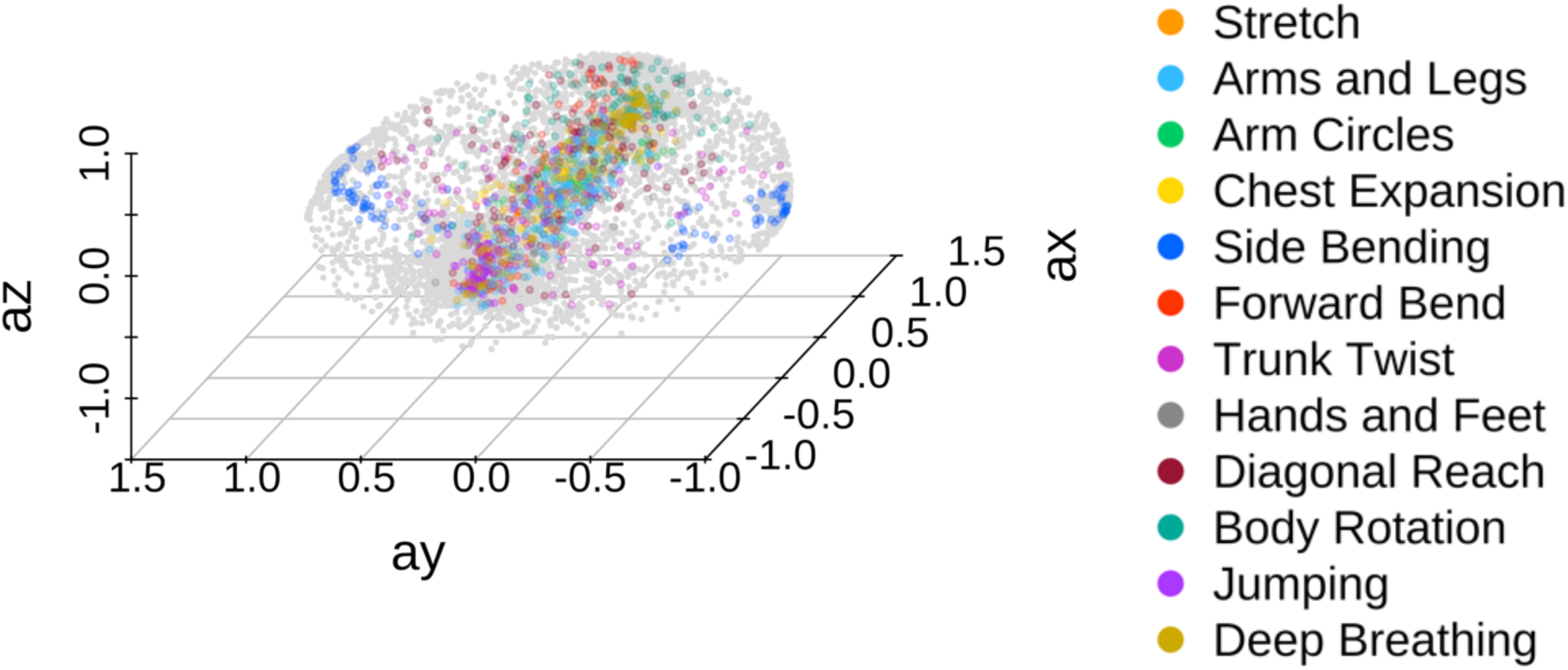
Bin-wise mean coordinates plotted in the MDS state space. The mean coordinate of each bin was superimposed on the frame-wise points (gray), with bin means color-coded according to exercise category. Although the same MDS state space as in Figure 2 was used, the viewing angle was changed to facilitate visualization of the spread along each axis.

### 3.2 Analysis of Distances in the MDS State Space

Next, statistical analyses were conducted to examine the effects of individual differences and exercise category on inter-point distances within the MDS state space. A two-way ANOVA was performed using an exercise factor, indicating whether the two points belonged to the same or different exercises, and a person factor, indicating whether the exercises were performed by the same or different participants. The results are summarized in Figure 4. Significant main effects of the exercise factor and person factor, as well as a significant interaction, were observed (exercise factor: *F*(1, 34) = 969.389, *p* < .001, 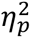 = .966; person factor: *F*(1, 34) = 17.062, *p* < .001, 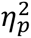 = .334; interaction: *F*(1, 34) = 16.765, *p* < .001, 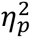 = .330).

**Figure 4.**
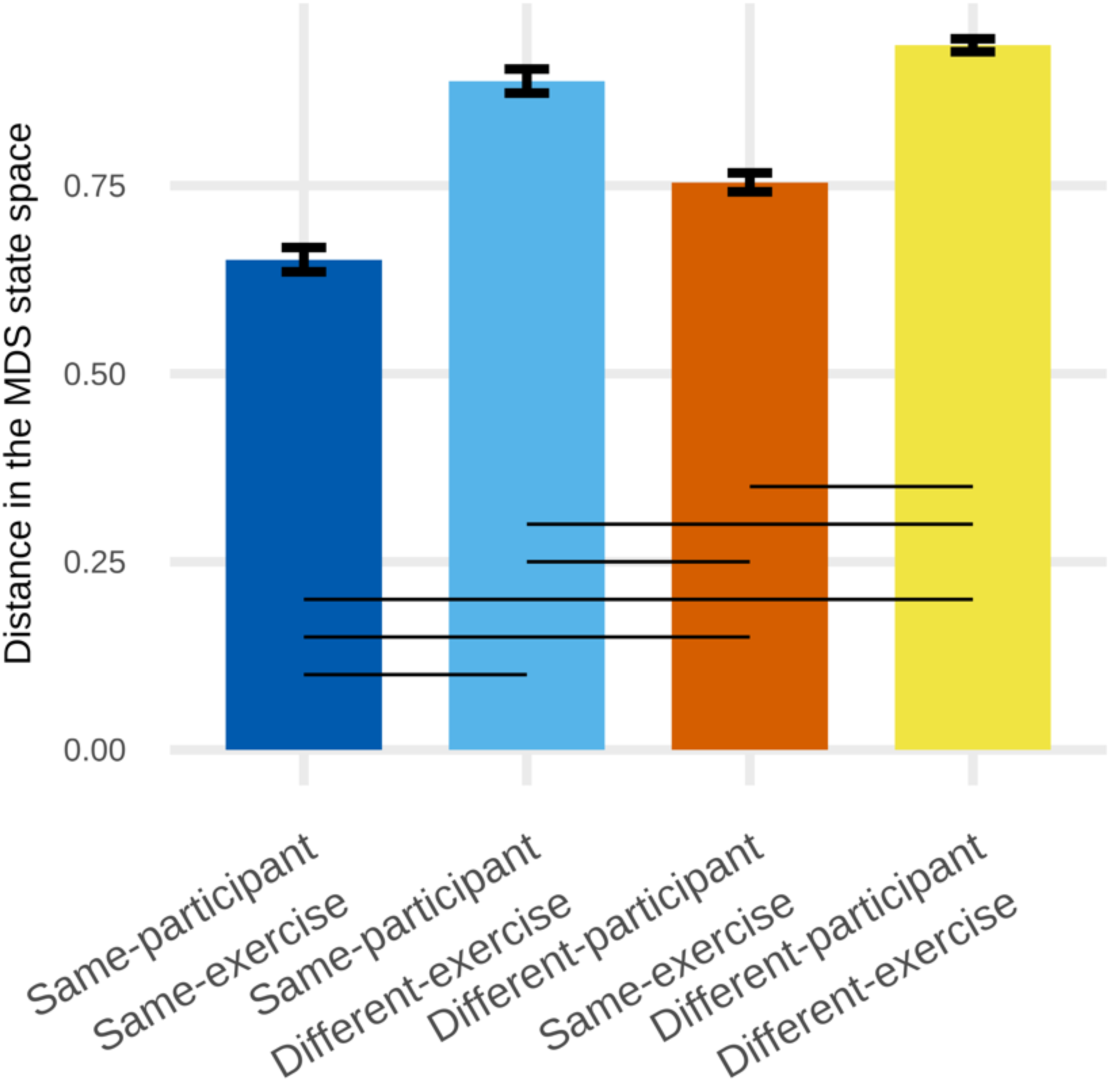
Distances in the MDS state space for the four conditions defined by participant and exercise factors. Mean distances for the four conditions, pooled across exercise categories. Error bars indicate standard errors. Horizontal lines below the graph indicate comparisons showing significant differences between connected conditions (*p* < .05).

Post hoc comparisons with Holm correction revealed significant differences among all four conditions. Distances were shortest when the same participant performed the same exercise, compared with the same participant performing different exercises (*t*(34) = -21.138, *p* < .001, *d* = -4.426), different participants performing the same exercise (*t*(34) = 4.717, *p* < .001, *d* = 1.916), and different participants performing different exercises (*t*(34) = -13.994, *p* < .001, *d* = -5.313). In contrast, distances were longest when different participants performed different exercises, compared with the same participants performing different exercises (*t*(34) = 2.851, *p* = .007, *d* = 0.887) and different participants performing the same exercises (*t*(34) = -24.459, *p* < .001, *d* = -3.397). Furthermore, distances were significantly shorter when the same exercise was performed by different participants than when different exercises were performed by the same participants (*t*(34) = -7.318, *p* < .001, *d* = -2.510).

Next, the effects of exercise category were examined across the 12 exercise categories (Figure 5). A three-way ANOVA was conducted by adding exercise category as a third factor to the two-factor model described above. Violations of the sphericity assumption were observed for the exercise-category factor and for the exercise category × exercise, exercise category × person, and exercise category × exercise × person interactions. Therefore, Greenhouse–Geisser corrections were applied. The analysis revealed significant main effects for all three factors and significant effects for all four interactions (exercise category: *F*(2.546, 86.570) = 139.647, *p* < .001, 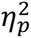 = .804; exercise factor: *F*(1.000, 34.000) = 1126.413, *p* < .001, 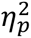 = .971; person factor: *F*(1.000, 34.000) = 13.601, *p* < .001, 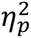 = .286; exercise category × person: *F*(2.546, 86.570) = 5.167, *p* = .004, 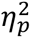= .132; exercise × person: *F*(1.000, 34.000) = 8.487, *p* = .006, 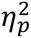 = .200; exercise category × exercise: *F*(2.496, 84.880) = 66.242, *p* < .001, 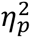 = .661; exercise category × exercise × person: *F*(2.496, 84.880) = 6.907, *p* < .001, 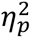 = .169).

**Figure 5.**
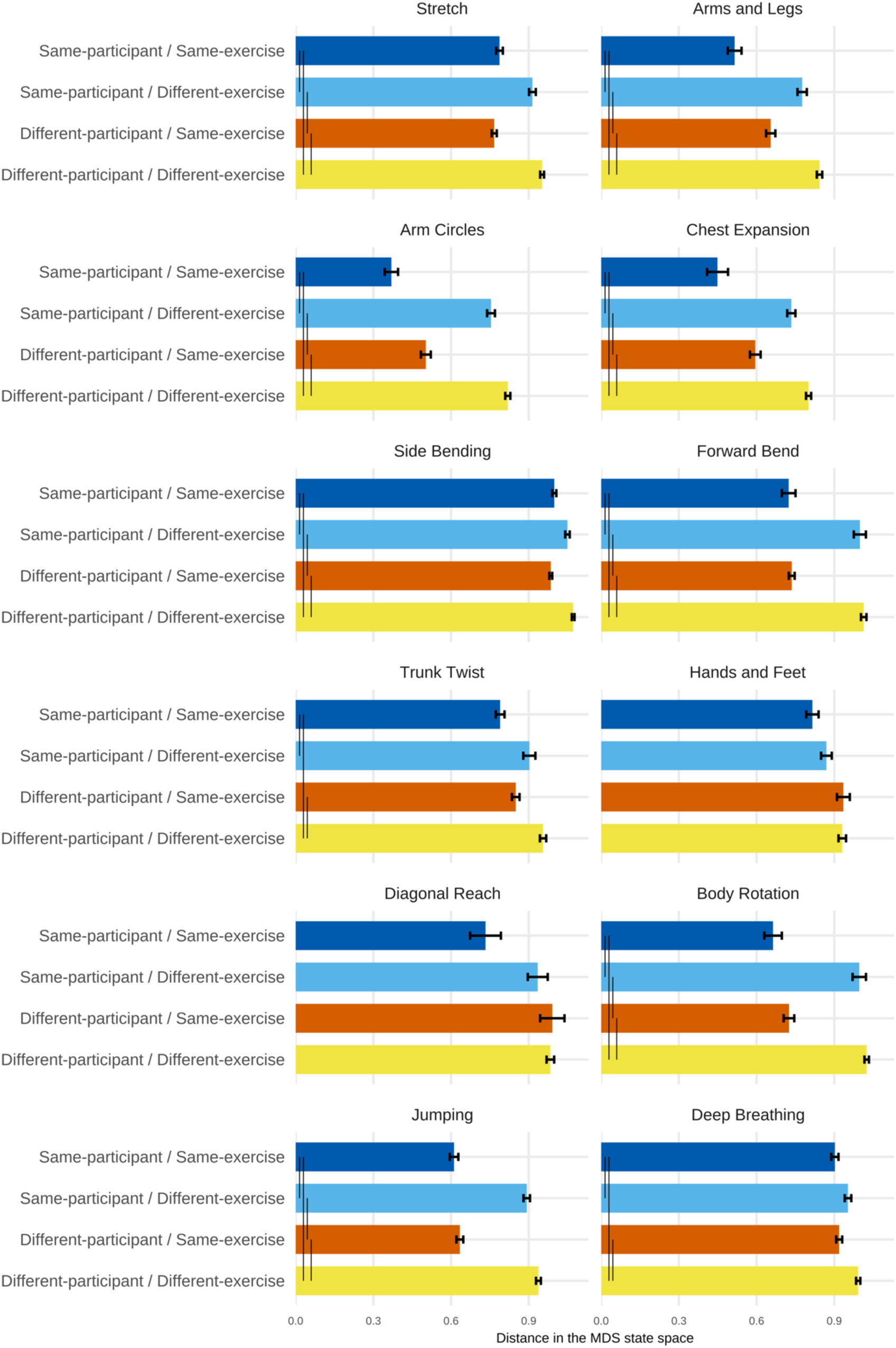
Mean distances in the MDS state space for each exercise category. Mean distances in the MDS state space for the four conditions defined by participant and exercise factors for each of the 12 exercise categories included in Radio Calisthenics. Error bars indicate standard errors. Because this study focused only on comparisons among the four conditions within each exercise, vertical lines within the bars indicate comparisons that were significant in the post hoc tests for the four-condition comparisons within each exercise.

Subsequent Holm-corrected simple-effects analyses were conducted to examine the three-way interaction. The present study focused specifically on comparisons among the four combinations of exercise factor (same vs. different exercise) and person factor (same vs. different participant) within each exercise category. The 12 exercise categories were classified into three broad patterns according to the locations of significant differences. The significant comparisons and corresponding statistics are summarized in Table 1 and Supplementary Table 1.

**Table 1.** Post hoc comparisons for the three-way interaction in the three-way ANOVA including exercise category. Only comparisons that remained significant after Holm correction are shown. SS, SD, DS, and DD represent the Same-participant/Same-exercise, Same-participant/Different-exercise, Different-participant/Same-exercise, and Different-participant/Different-exercise conditions, respectively.

| Pattern | Exercises | Significant Comparisons |
| --- | --- | --- |
| 1 | Stretch, Arms and Legs, Arm Circles, Chest Expansion, Side Bending, Forward Bend, Body Rotation, Jumping | SS < SD |
|  |  | SS < DD |
|  |  | DS < SD |
|  |  | DS < DD |
| 2 | Trunk Twist, Deep Breathing | SS < SD |
|  |  | SS < DD |
|  |  | DS < DD |
| 3 | Hands and Feet, Diagonal Reach. | No Significant Differences |

The first pattern was observed for Stretch, Arms and Legs, Arm Circles, Chest Expansion, Side Bending, Forward Bend, Body Rotation, and Jumping. In these exercises, distances were significantly shorter for Same exercise comparisons than for Different exercise comparisons, both within the Same participant and Different participant conditions.

The second pattern was observed for Trunk Twist and Deep Breathing. In these exercises, distances were significantly shorter for same-exercise comparisons than for Different exercise comparisons within both the Same participant and Different participant conditions. In addition, distances in the Same-participant/Same-exercise condition were significantly shorter than those in the Different-participant/Different-exercise condition.

Finally, no significant differences were observed among any of the four conditions for Hands and Feet and Diagonal Reach.

Because the patterns of distances across the four conditions differed among exercise categories, we explored factors that might account for these differences. As representative examples of Patterns 1 and 2, Arm Circles and Deep Breathing were selected for further examination. The primary difference between these two patterns was whether a significant difference was observed between the Same-participant/Different-exercise condition and the Different-participant/Same-exercise condition. In addition, Deep Breathing showed a distinctive characteristic in that the distance observed in the Same-participant/Same-exercise condition appeared comparable to the distance observed in the Different-participant/Different-exercise condition for many other exercises.

Because Deep Breathing was performed more slowly than the other exercises, within-bin variability was examined (Figure 6). Within-bin variability was quantified as the mean distance between the centroid of each bin and the coordinates of the individual frames contained within that bin. As a result, Deep Breathing exhibited approximately half the within-bin variability of Arm Circles. However, several other exercises showed a similar level of variability. Forward Bend, Diagonal Reach, and Body Rotation exhibited within-bin variability comparable to that of Deep Breathing and lower than that observed for most other exercises. In contrast, Arms and Legs and Chest Expansion, like Arm Circles, exhibited relatively large within-bin variability.

**Figure 6.**
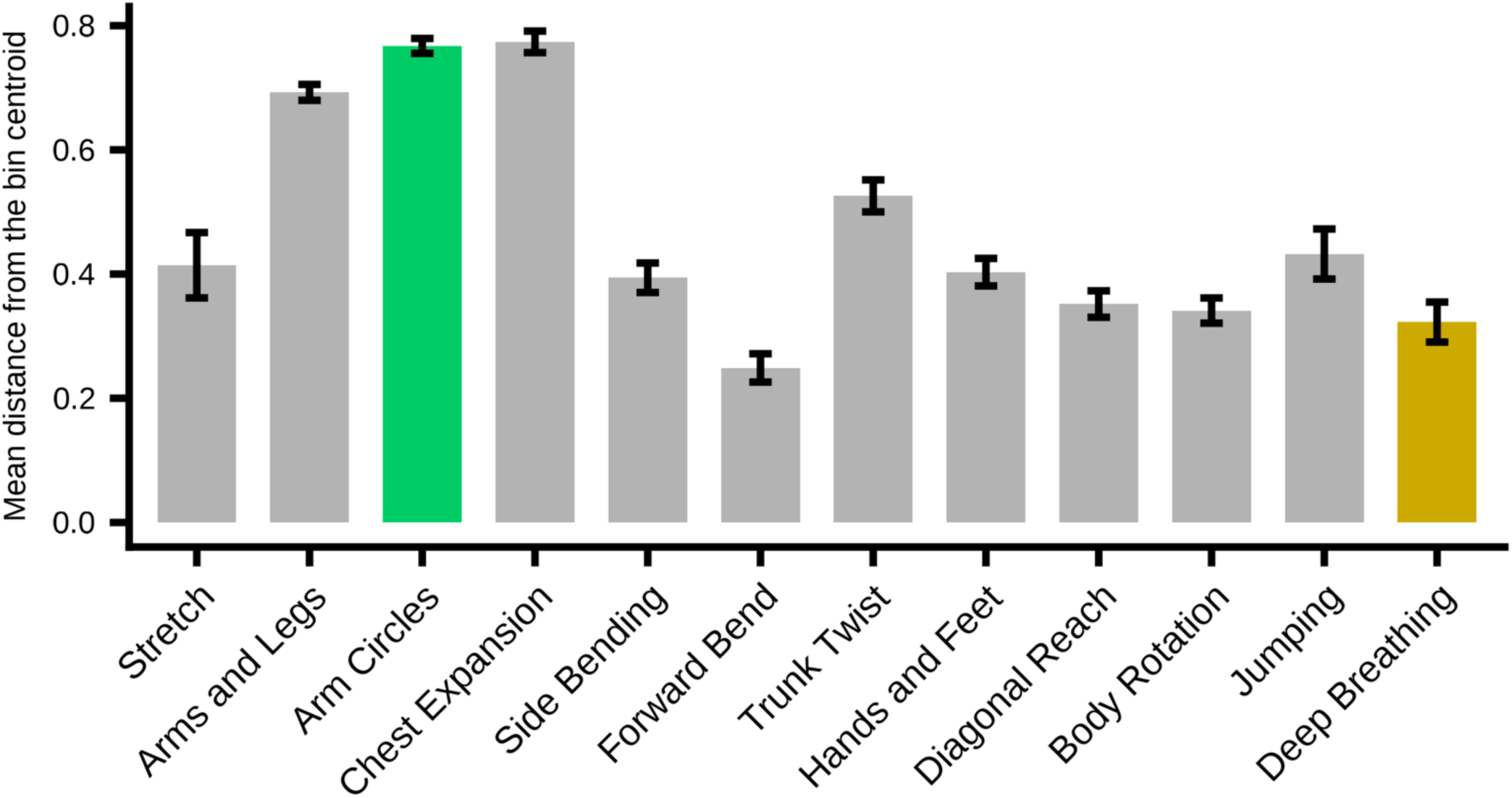
Within-bin variability for each exercise in Radio Calisthenics. For each frame, the distance from the bin centroid of the corresponding bin was calculated and averaged for each exercise. Error bars indicate standard errors.

Finally, to examine whether differences between motion capture recording sessions were reflected in the MDS state space, we performed a three-way ANOVA in which the exercise-category factor was replaced with a recording-session factor (same day vs. different day) (Figure 7). The main effect of recording session was not significant, *F*(1, 32) = 2.941, *p* = .096, 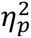 = .084. We further examined the Holm-adjusted post hoc comparisons to determine whether the four combinations of participant and exercise conditions differed between the same-day and different-day recording sessions. No significant differences were found for any of the four conditions.

**Figure 7.**
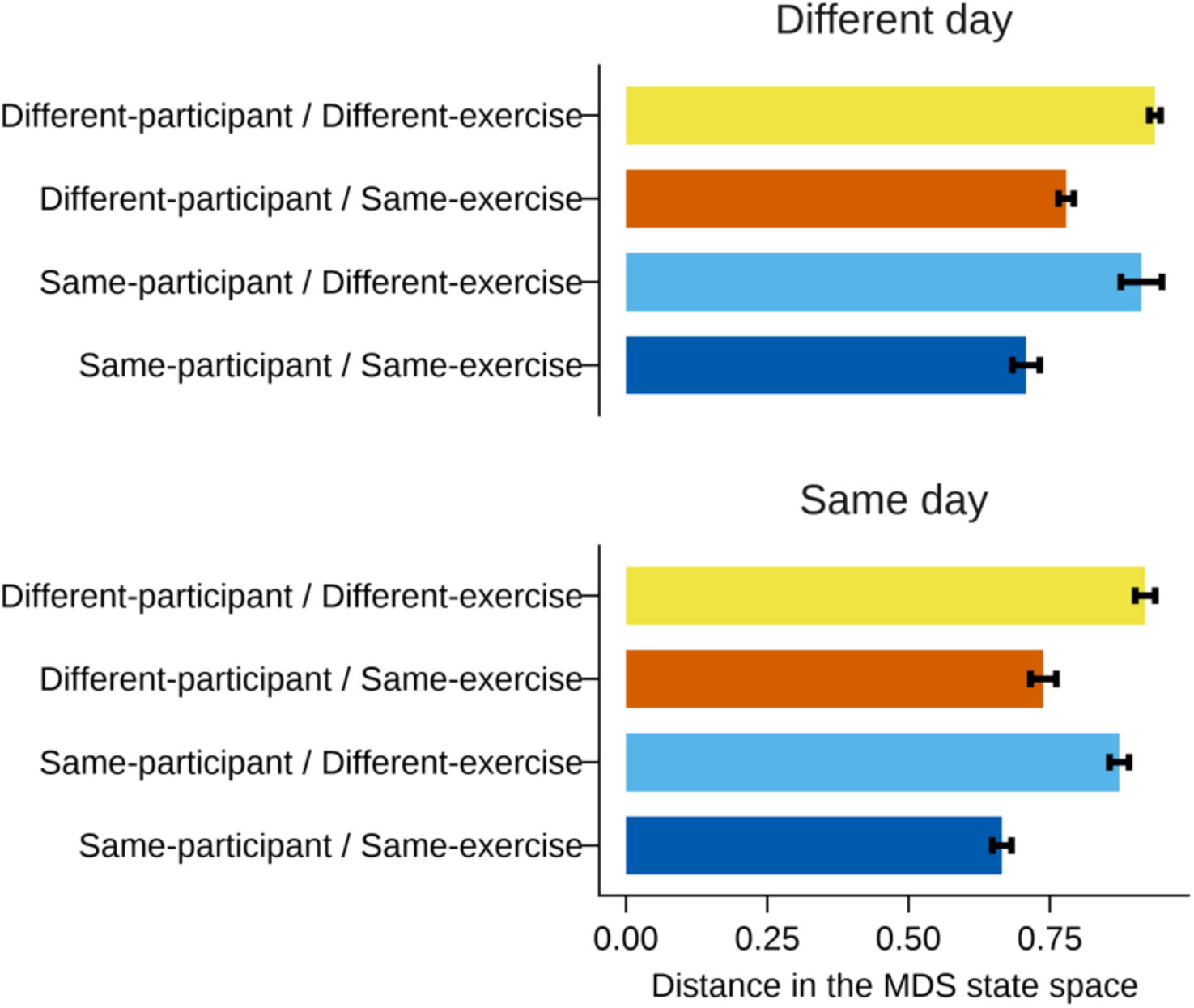
Distances between points in the MDS state space when considering recording day. Distances were summarized according to whether recordings were obtained on the same or different recording days. Error bars indicate standard errors.

## 4. Discussion

The present study examined the validity of MDS as a method for representing and summarizing whole-body movement in an interpretable low-dimensional space. Using Radio Calisthenics as an example of whole-body movement, we focused on whether the dimensionality reduction produced an interpretable representation that reflected exercise categories, which is a prerequisite for future investigations of its relationship with impression evaluation. Specifically, we evaluated validity according to two broad criteria: (1) similarities between postures should be reflected in inter-point distances within the state space, without points collapsing into a central cluster or being arranged at uniform intervals; and (2) points representing the same exercise should be located closer together regardless of individual differences or differences in body position and orientation. The first criterion was intended to verify that the MDS procedure behaved as expected. Because MDS determines inter-point distances based on similarities among data points, it was first necessary to confirm whether the different exercises included in Radio Calisthenics could be represented as distinguishable distributions within the MDS state space. The second criterion was intended to test whether exercise category, rather than individual differences, would primarily determine inter-point distances within the MDS state space. Specifically, we examined whether points would cluster according to exercise category rather than participant, with points representing the same exercise located closer together and points representing different exercises located farther apart.

The resulting MDS state space showed that the plotted points did not collapse around the center, but instead formed a distribution resembling the surface of a sphere (Figure 2b). In addition, some regions of the state space were relatively dense whereas others were sparse, and the points were not arranged at uniform intervals. These observations were consistent with the first criterion.

The correspondence between exercise category and movement similarity represented in the state space was examined by performing ANOVAs on the inter-point distances between points plotted in the state space. A two-way ANOVA was conducted with a person factor, indicating whether the exercises were performed by the same participant, and an exercise factor, indicating whether the exercises belonged to the same exercise category. Significant differences were observed among all four conditions. Distances were shortest when the same participant performed the same exercise and longest when different participants performed different exercises. In addition, when comparing exercises performed by the same participant, distances were longer for different exercises than for the same exercise. These results indicate that exercise category was reflected in distances within the state space more strongly than whether the exercises were performed by the same participant. Combined with the observation in Figure 2b that points from different trials and participants overlapped with one another, these findings suggest that the second criterion was also generally satisfied: the representation did not form participant-specific clusters, and exercise category was reflected in the distances more strongly than individual differences. Thus, the validity criteria proposed in the present study were generally met, indicating that dimensionality reduction using an MDS state space has potential utility as a method for summarizing whole-body movement. In addition, Figure 3 suggested that some exercise categories tended to spread along different dimensions of the state space, indicating that certain exercise characteristics may be reflected in specific dimensions.

Nevertheless, the validity of dimensionality reduction should be considered from perspectives beyond the criteria examined in the present study. Ideally, the analysis should include the full range of postures that humans can assume. In the present study, however, we focused only on a limited set of movements included in Radio Calisthenics. We therefore also conducted a three-way ANOVA to examine whether the patterns of distances within the state space differed across exercise categories. The most common pattern, observed in 8 of the 12 exercise categories, was that distances were shorter for same-exercise comparisons than for different-exercise comparisons within both the same-participant and different-participant conditions, and that the Same-participant/Same-exercise condition showed shorter distances than the Different-participant/Different-exercise condition. In addition, no significant difference was observed between the Same-participant/Same-exercise and Different-participant/Same-exercise conditions, suggesting that the same exercises tended to be represented by similar distances regardless of the performer. Overall, these findings indicate that Same-exercise comparisons tended to yield shorter distances than Different-exercise comparisons. However, the remaining four exercise categories showed different patterns. In particular, for Deep Breathing, the distance in the Same-participant/Same-exercise condition appeared comparable to the distance in the Different-participant/Different-exercise condition for other exercises.

To explore why the patterns of significant differences varied across exercise categories, we focused on Deep Breathing, which showed a distinctive pattern compared with the other exercises. Deep Breathing is performed at the end of Radio Calisthenics and does not require a transition to a subsequent exercise. It was also performed more slowly than the other exercises. Therefore, we focused on differences in movement speed and compared within-bin variability. Consistent with the prediction that slower exercises would show smaller within-bin variability, Deep Breathing showed approximately half the within-bin variability of Arm Circles, which was considered a representative example of Pattern 1 and showed particularly short distances in the Same-participant/Same-exercise condition. Although within-bin variability may be related to distances in the state space, several other exercises also showed within-bin variability as small as that of Deep Breathing. These exercises did not necessarily show longer distances in the Same-participant/Same-exercise condition; some showed the same pattern of significant distance differences as Arm Circles. Thus, small within-bin variability alone could not fully explain the differences in distances within the state space. Exercise characteristics specific to Deep Breathing may have been related to this pattern.

Arm Circles and Deep Breathing both involve large arm movements following circular trajectories. However, the two exercises may differ in the range of interpretations and executions allowed by the demonstration. In Arm Circles, the hands are crossed in front of the body, and both arms are then rotated alternately outward and inward in time with the counts in the video. Given the overall tempo of the music, this exercise is generally performed with a certain degree of momentum, making it difficult to vary movement speed during execution. Because the arms must complete a full rotation while extended, and because the exercise is performed at a relatively constant speed, participants tended to follow similar trajectories across repetitions within the same participant. In contrast, Deep Breathing is slower than the other exercises, and because no subsequent exercise follows it, participants could adjust the speed of their arm movements from moment to moment. In addition, because the time spent with the arms raised was relatively long, individual interpretation may have influenced how high the arms were raised. As a result, even across repetitions of Deep Breathing, the timing and height of the arm movement may have varied more easily, resulting in greater between-repetition variability within participants compared with other exercises. In the present experiment, participants were instructed to perform the sequences while watching the demonstration without overexerting themselves, and no scoring or feedback was provided regarding how closely they followed the demonstration. Therefore, the degree to which participants adhered to the demonstration was left to each individual. Radio Calisthenics is incorporated into school education in Japan and is also practiced in community activities. Because the experiment assumed that all participants were already familiar with the basic sequences, no prior learning session or practice trials were conducted before recording. Given that prior experience with Radio Calisthenics was not controlled, the task and experimental context may have allowed individual movement habits to emerge, depending on the exercise category.

As an additional check, we examined the effect of recording day. A three-way ANOVA was conducted by adding a recording-day factor to the exercise and person factors. The main effect of recording day was not significant, indicating that inter-point distances did not differ significantly depending on whether the data were recorded on the same day. Because of differences in camera connectivity, recordings obtained on the same day were sometimes processed using calibrations based on different numbers of cameras. Therefore, being recorded on the same day was not equivalent to sharing the same calibration data. Nevertheless, the present analyses did not reveal clear effects of repeated recordings by the same participant or of whether recordings were obtained on the same day.

A future challenge is to quantitatively relate movement features to the configuration of the state space in order to examine the relationship between summarized whole-body movement and impression evaluation. It will also be necessary to clarify what movement features are represented by each dimension of the state space. The exercise-category analyses suggested that spatiotemporal features, such as movement speed, may influence condition-specific distance patterns within the state space. Possible contributing factors include within-bin variability and differences in the degree of interpretability of the demonstrated movements, but their quantitative relationships remain unclear. To identify the factors that determine distance patterns within the state space, future studies should examine individual differences in movement trajectories and execution timing using more rigorous measurement and analysis methods with larger samples. In addition, when the points in the state space were color-coded by exercise category (Figure 3), some exercises appeared to correspond to particular dimensions of the state space. Because the orientation of MDS solutions can rotate depending on the analysis, the first to third dimensions observed in the present analysis do not necessarily correspond to fixed movement features. Nevertheless, the relationships among movements distributed along each dimension may contain independent components. Even within the limited movement repertoire of Radio Calisthenics, Side Bending, which involves lateral flexion of the upper body, tended to extend along a dimension different from that of other movements primarily involving the arms. This suggests that movements involving substantially different body regions may be represented as differences along distinct dimensions. Clarifying the relationships among movements based on the overall structure of the state space may deepen our understanding of which forms of inter-body-part coordination are most readily expressed in whole-body movement.

Relatedly, the present study has two main limitations. First, the movements examined in this study were limited to a restricted repertoire of sequential movements. Radio Calisthenics was selected because it involves whole-body movement and is familiar to many Japanese people. However, many of the movements included in Radio Calisthenics primarily involve the arms and do not include large leg movements or substantial changes in whole-body posture. Therefore, the state space constructed in the present study was likely biased toward upper-body movements, particularly arm movements. To investigate the relationship between whole-body movement and impression evaluation, as well as to promote a broader understanding of whole-body movement itself, it will be necessary to construct a state space that includes a wider variety of whole-body movements that humans can perform. One possible future approach is to expand the range of movements included in the state space by using large-scale motion datasets.

The second limitation is the small number of participants. Because the present study targeted participants who already knew Radio Calisthenics and did not require practice, the range of movement skill levels was limited. Some forms of movement evaluation, such as aesthetic evaluation, are strongly related to skill level. Further work is needed to examine whether similar patterns would be observed when movements are performed by participants encountering the movements for the first time, or conversely, by participants highly skilled in the movements, such as the models in the demonstration video. It will also be important to examine whether differences in skill level are reflected as individual differences in the state-space representation.

## 5. Conclusion

In the present study, we evaluated whether the state space generated by MDS provides a valid representation for summarizing whole-body movement. Specifically, we examined whether the state space consistently represented movement similarity by reflecting exercise category more strongly than individual differences. Using Radio Calisthenics data collected from five participants, distances between points in the MDS state space were compared across four conditions defined by whether the exercises were performed by the same participant and whether they belonged to the same exercise category.

The results showed that exercise category was reflected in the distances between points more strongly than participant identity. Distances were shorter between the same types of exercises and longer between different types of exercises. These findings suggest that whole-body movements can be represented in a low-dimensional space such that points representing the same exercise categories are located closer together. However, this tendency was more evident for some exercise categories than for others. Furthermore, for some exercises, the distance under the Same-participant/Same-exercise condition was comparable to that under the Different-participant/Different-exercise condition for other exercises. These findings also suggest that the distance relationships varied across exercise categories. Such variation may be associated not only with within-bin variability but also with exercise-specific characteristics, including the degree of freedom with which participants interpreted the reference movement. Future studies should include a more diverse range of participants with different levels of movement proficiency and extend the analysis to a wider variety of whole-body movements beyond Radio Calisthenics in order to further examine the generalizability and validity of the MDS state space.

## Acknowledgments

MW, AK, and TA were supported by the JST Moonshot R&D Program, Grant Number JPMJMS2291, Japan.

## Author Contributions

MW conceived the study; collected the data; performed the formal analyses; developed the methodology; conducted the investigation; managed the project; and wrote the original draft of the manuscript. AK contributed to data collection, assisted with the experimental setup, and provided advice on the experimental procedures. TA supervised the study, contributed to the study design, and provided guidance on the analytical methods.

All authors reviewed the manuscript, approved the final version, and agreed to its publication.

Generative AI was used to assist with code development for statistical analyses and figure preparation, as well as language editing. All generated content was reviewed, verified, and revised by the authors, who take full responsibility for the final manuscript.

