## Supplementary Information for "Validity of Low-Dimensional Representation of Whole-Body Movements"

**Supplemental information**

Supplementary Table 1. Results of the post hoc multiple comparisons for the three-way interaction.

SS = same participant performing the same exercise; SD = same participant performing different exercises; DS = different participants performing the same exercise; DD = different participants performing different exercises.

| Pattern | Exercise | Condition1 | Condition2 | df | t | p | Cohen's d |
|---|---|---|---|---|---|---|---|
| 1 | Stretch | DS | SS | 34 | -1.193 | 1.000 | -0.247 |
| | | | DD | 34 | -18.643 | < .001 | -2.244 |
| | | | SD | 34 | -9.716 | < .001 | -1.791 |
| | | SS | DD | 34 | -10.111 | < .001 | -1.997 |
| | | | SD | 34 | -8.504 | < .001 | -1.543 |
| | | DD | SD | 34 | 2.618 | 1.000 | 0.454 |
| | Arms and Legs | DS | SS | 34 | 4.295 | 0.052 | 1.690 |
| | | | DD | 34 | -17.307 | < .001 | -2.289 |
| | | | SD | 34 | -4.983 | 0.008 | -1.475 |
| | | SS | DD | 34 | -11.242 | < .001 | -3.980 |
| | | | SD | 34 | -15.873 | < .001 | -3.165 |
| | | DD | SD | 34 | 3.383 | 0.550 | 0.814 |
| | Arm Circles | DS | SS | 34 | 3.951 | 0.129 | 1.615 |
| | | | DD | 34 | -28.618 | < .001 | -3.842 |
| | | | SD | 34 | -10.526 | < .001 | -3.055 |
| | | SS | DD | 34 | -15.098 | < .001 | -5.457 |
| | | | SD | 34 | -23.078 | < .001 | -4.671 |
| | | DD | SD | 34 | 3.600 | 0.321 | 0.787 |
| | Chest Expansion | DS | SS | 34 | 3.551 | 0.362 | 1.769 |
| | | | DD | 34 | -12.699 | < .001 | -2.499 |
| | | | SD | 34 | -5.093 | 0.006 | -1.690 |
| | | SS | DD | 34 | -9.857 | < .001 | -4.268 |
| | | | SD | 34 | -11.660 | < .001 | -3.459 |
| | | DD | SD | 34 | 3.645 | 0.285 | 0.809 |
| | Side Bending | DS | SS | 34 | -1.360 | 1.000 | -0.164 |
| | | | DD | 34 | -14.107 | < .001 | -1.049 |

| | | | | | | | |
|---|---|---|---|---|---|---|---|
| | | | SD | 34 | -6.552 | < .001 | -0.781 |
| | | SS | DD | 34 | -7.379 | < .001 | -0.885 |
| | | | SD | 34 | -5.511 | 0.002 | -0.618 |
| | | DD | SD | 34 | 2.251 | 1.000 | 0.267 |
| | Forward Bend | DS | SS | 34 | 0.487 | 1.000 | 0.145 |
| | | | DD | 34 | -16.922 | < .001 | -3.374 |
| | | | SD | 34 | -11.494 | < .001 | -3.193 |
| | | SS | DD | 34 | -12.197 | < .001 | -3.518 |
| | | | SD | 34 | -11.104 | < .001 | -3.337 |
| | | DD | SD | 34 | 0.673 | 1.000 | 0.181 |
| | Body Rotation | DS | SS | 34 | 1.604 | 1.000 | 0.746 |
| | | | DD | 34 | -13.278 | < .001 | -3.644 |
| | | | SD | 34 | -9.813 | < .001 | -3.296 |
| | | SS | DD | 34 | -10.622 | < .001 | -4.390 |
| | | | SD | 34 | -9.771 | < .001 | -4.042 |
| | | DD | SD | 34 | 1.339 | 1.000 | 0.347 |
| | Jumping | DS | SS | 34 | 0.982 | 1.000 | 0.273 |
| | | | DD | 34 | -39.701 | < .001 | -3.684 |
| | | | SD | 34 | -13.843 | < .001 | -3.136 |
| | | SS | DD | 34 | -15.423 | < .001 | -3.957 |
| | | | SD | 34 | -24.369 | < .001 | -3.409 |
| | | DD | SD | 34 | 2.742 | 1.000 | 0.548 |
| 2 | Trunk Twist | DS | SS | 34 | 2.420 | 1.000 | 0.734 |
| | | | DD | 34 | -10.347 | < .001 | -1.281 |
| | | | SD | 34 | -2.165 | 1.000 | -0.638 |
| | | SS | DD | 34 | -6.730 | < .001 | -2.015 |
| | | | SD | 34 | -7.347 | < .001 | -1.371 |
| | | DD | SD | 34 | 2.214 | 1.000 | 0.643 |
| | Deep Breathing | DS | SS | 34 | 0.847 | 1.000 | 0.204 |
| | | | DD | 34 | -10.328 | < .001 | -0.892 |
| | | | SD | 34 | -2.054 | 1.000 | -0.413 |
| | | SS | DD | 34 | -4.887 | 0.010 | -1.096 |
| | | | SD | 34 | -4.741 | 0.015 | -0.617 |

| | | | | | | | |
|---|---|---|---|---|---|---|---|
| | | DD | SD | 34 | 2.644 | 1.000 | 0.479 |
| 3 | Hands and Feet | DS | SS | 34 | 2.923 | 1.000 | 1.454 |
| | | | DD | 34 | 0.339 | 1.000 | 0.052 |
| | | | SD | 34 | 2.090 | 1.000 | 0.794 |
| | | SS | DD | 34 | -3.119 | 1.000 | -1.402 |
| | | | SD | 34 | -2.873 | 1.000 | -0.659 |
| | | DD | SD | 34 | 2.359 | 1.000 | 0.743 |
| | Diagonal Reach | DS | SS | 34 | 3.172 | 0.901 | 3.131 |
| | | | DD | 34 | 0.216 | 1.000 | 0.091 |
| | | | SD | 34 | 1.061 | 1.000 | 0.682 |
| | | SS | DD | 34 | -3.564 | 0.352 | -3.040 |
| | | | SD | 34 | -3.850 | 0.168 | -2.449 |
| | | DD | SD | 34 | 1.451 | 1.000 | 0.591 |
